# Age-related balance difficulties during shoulder checks increase cycling risk in older adults

**DOI:** 10.1101/2022.03.01.481993

**Authors:** Maarten Afschrift, Frea Deroost, Anouck Matthijs, Theresa De Ryck, Friedl De Groote, Jean-Jacques Orban De Xivry

## Abstract

Older adults are more often involved in bike accidents than other age groups, especially in single-sided accidents (crashes without another road user). This suggests that older adults have more difficulty maintaining balance while cycling, likely due to age-related sensorimotor decline. A particularly challenging maneuver in traffic is the shoulder check (turning the upper body to look behind while maintaining balance and direction). This study investigates how age-related sensorimotor deficits impact this skill.

40 young (22.86 ± 1.53 years) and 41 older participants (62.73 ± 1.57 years) cycled in a straight lane while performing a shoulder check to identify the color of an object behind them. We recorded the task-errors (interrupted cycling, incorrect color identification, cycling outside the lane) and computed the steering angle, rotation of the frame, pelvis and torso, and task duration from inertial measurement unit data.

Older adults made more task-errors than young participants; one-third failed the task due to mistakes such as misidentifying the color, losing balance, or leaving the lane. In the successful trials, older adults showed greater steering variability, increased pelvic rotation relative to the frame, and took longer to complete the task than the young participants.

These findings suggest that age-related difficulties in maintaining balance while head-turning may contribute to the higher rate of single-sided bike accidents among older cyclists. Infrastructure adaptations (e.g., wider bike lanes) and individual-level interventions (e.g., mirrors) that alleviate the difficulties older adults experience when performing a shoulder check could improve safety for people from this age group.

## 1. Introduction

Cycling is a common mode of transport and a popular recreational activity for all age groups across multiple countries (e.g. Belgium, the Netherlands, Denmark), valued for its sustainable, social, and health-related benefits. In Flanders (Belgium), 83% of the adult population uses a bicycle (VAB fietsbarometer 2018). This popularity extends to older adults, as one in three older adults uses a bicycle at least once a year [1,2]. In addition, the overall number of cyclists continues to rise annually, including older adults [3].

Despite the benefits of cycling, cycling safety remains a major concern. As bicycle use increases, so too does the incidence of cycling-related accidents. In 2023, a record number of 11,595 cyclists were injured in traffic accidents in Flanders [4]. Older adults are disproportionately affected by these incidents, not only being more frequently involved in accidents compared to younger cyclists but also experiencing more severe consequences [5]. Causes of traffic accidents during cycling are related to the interaction between (1) the human on the bicycle, (2) the mechanics of the bicycle, and (3) the environment (including other road users) [6–8]. Accident statistics suggest that these interactions differ in different subpopulations. In young or inexperienced cyclists, most cycling accidents occur in interaction with motorized vehicles, whereas in older adults, there is a strong increase in the number of cyclists that are admitted to the hospital due to single-sided bicycle accidents (i.e., accidents involving the cyclist only) [9]. Between 60% and 95% of cyclists admitted to hospitals are the result of single-sided bicycle accidents [10]. There is a 37% increase in single-sided accidents in older adults compared to the average of all cyclists and a 24% increase in collisions with another obstacle than a road user [2].

The increase in single-sided accident rate in older subjects suggests that older adults have more difficulty controlling cycling direction and balance while bicycling in traffic. Bulsink et al. [11] suggested that older adults need more effort to restore from perturbations based on the observation that older subjects use additional strategies, such as outward knee movement, to control balance. Neuro-physiological changes in the human body with aging [12], such as frailty, slower reaction times [13], less accurate sensory abilities [14,15], reduced range of motion in the musculoskeletal system [16] and higher mental workload experienced [17,18], might be associated with the observed differences in balance control strategies [11] and the increased number of single sided accidents. For example, the increase of accident risk in older cyclists (on an e-bike) has been related to a relatively higher mental workload for older cyclists (65+ years) compared to a group of middle-aged cyclists (30-45 years) [18].

A particularly challenging scenario for older cyclists involves making a turn at an intersection or crossing the road, maneuvers that often results in single-sided accidents involving older adults [19]. This action requires the cyclist to perform a shoulder check to assess whether traffic is approaching from behind and if it is safe to turn, which involves rotating the upper body while maintaining balance and cycling direction. Age-related neurophysiological changes are believed to make this coordination especially difficult for older adults. Kovácsová et al. [20] demonstrated that a shoulder check movement combined with raising the hand for a left turn caused increased variability in steering angle for older compared to middle-age groups. In addition, this was associated with a lower ability to identify the number of hand raised by the experimenter behind the cyclists. This clearly shows that older people have limited abilities to remain stable on their bike while turning their upper body to check on potential road users. Yet, it is currently unknown which age-related sensorimotor deficits influence this ability.

To fill this gap, we instructed a group of young and a group of older participants to bike in a straight line while performing a shoulder check movement in order to identify the color of an object presented behind them. This task involves both the rotation of trunk and head for perception of the environment and the concurrent control of balance and cycling direction. Therefore, it allows us to investigate the ability of older adults to perform these two concurrent motor tasks. This task is ecologically valid as it is relevant for multiple traffic situations: such as identifying cars behind the cyclist when changing lane or overtaking another road user. We evaluated if the young and older adults could perform the task without errors (e.g. wrong identification, cyclist outside the test track, loss of balance) and if they performed steering corrections to maintain cycling direction.

## 2. Methods

### 2.1 participants

A group of 40 young (22.86 ± 1.53 years) and 41 older participants (62.73 ± 1.57 years, minimum 60 years and maximum 65 years) subjects participated in this study. Participants were recruited through mailings to local organizations and sport clubs for seniors, through local advertisements and by contacting the social network of the research team. Only participants who cycled at least once a month were included. The study was approved by the local ethics committee (SMEC G-2020-2410) and all participants provided their written informed consent.

### 2.2 Experimental procedure

We instructed participants to cycle on a self-designed test track that mimicked several real-world conditions on an aluminum bike with (e-bike) or without a motor (conventional bike) (order of the bike was randomized across participants, mass conventional bike = 19.8 kg, details in appendix). Participants biked on the track under three different conditions: first at self-selected speed, then twice more slowly than the self-selected speed and finally at self-selected speed while performing a concurrent dual-task. For this paper, we focus on the self-selected speed condition on the conventional bike but the results are identical for the e-bike and for the slow condition (Appendix 2). The subjects familiarized themselves with the cycling track for five minutes (two repetitions) with both the e-bike and the normal bike and subsequently performed one trial that was recorded and analyzed. In the present study we analyzed the part in the test track where subjects had to identify a yellow or red cone behind the cyclist by first looking over the left and secondly over the right shoulder (Figure 1). During the test, the experimenter wrote down (a) whether the participants lost balance and needed to place a foot on the ground or not, (b) whether the participants made mistakes in identifying the cone, (c) and whether they cycled outside the narrow lane with yellow cones (Figure 1).

**Figure 1:**
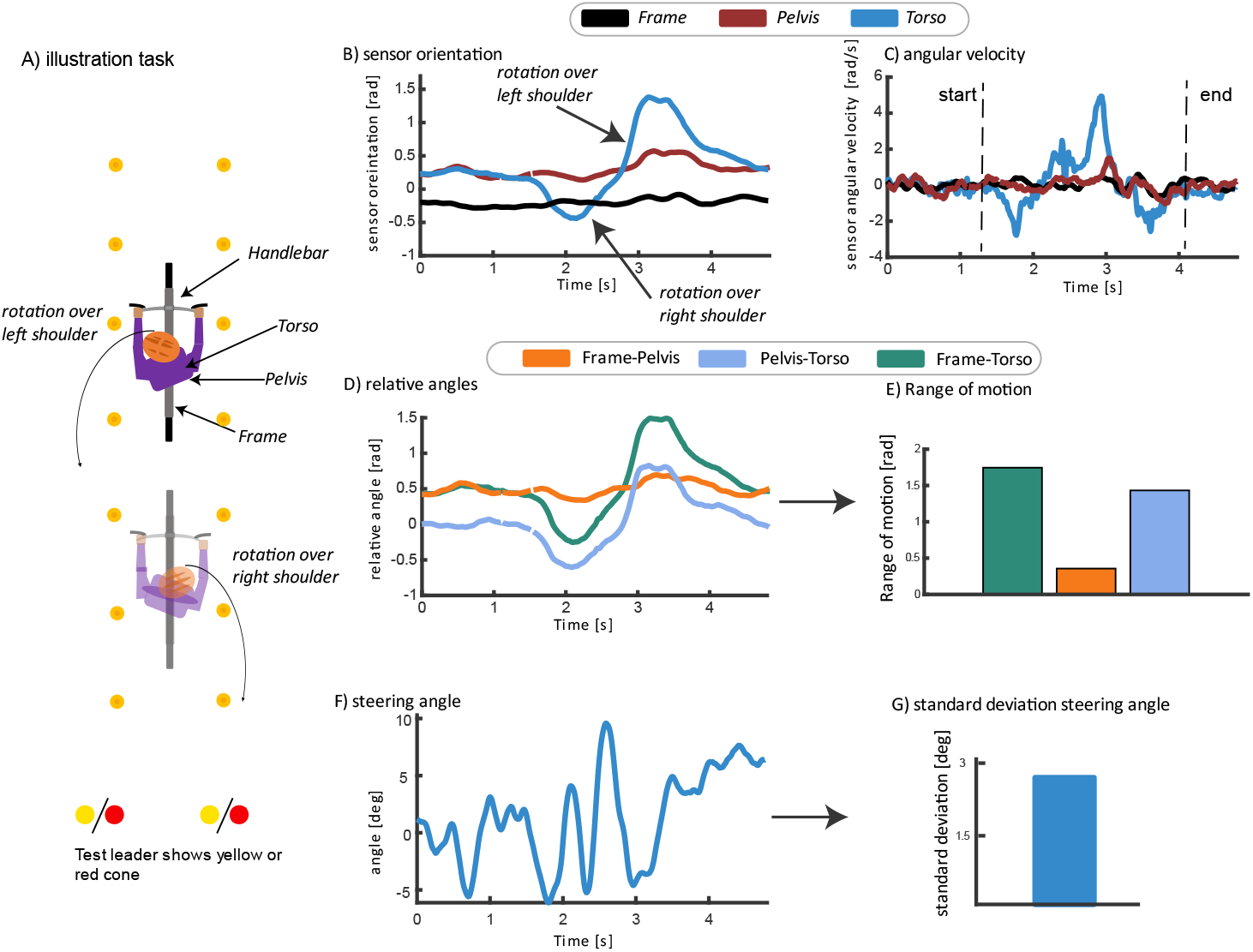
Overview of the kinematic analysis. Young and older subjects identified a yellow or red cone behind the cyclist by first looking over the left and secondly over the right shoulder while cycling in a narrow track between cones (A). The kinematics of the bike and cyclist were analyzed using IMU’s on the handlebar, frame, pelvis, torso. Sensor orientations were computed from the IMU raw data and orientations were evaluated around the vertical axis (B). The start and end of the rotation over the right and left shoulder was identified manually based on the angular velocity of sensors (C). Relative changes in orientation between the sensor on the frame, pelvis and torso were computed in the identified time-window to quantify the relative rotation of the segments (D). The range of motion was computed from the relative rotation between the sensors (E). The steering angle was computed as the orientation of the handlebars with respect to the frame along the axis of the stem (F), and the resulting standard deviation in steering angle was compared between the young and older subjects (G).

### 2.3 Kinematic analysis + determine steering angle

The kinematics of the cyclist and bike was analyzed using inertial measurement units (IMU) mounted on the handlebars, frame, pelvis and torso (MTw Awinda, Xsens, The Netherlands). IMU data from the handlebars and frame were collected at 100 Hz, while data from the pelvis and torso were recorded at 75 Hz. Sensor orientations were computed from the raw IMU data using the Xsens sensor fusion algorithm in proprietary software MT manager (Xsens, The Netherlands). Sensor orientations were exported as rotation matrices for further analysis.

The steering angle is defined as the rotation of the handlebars with respect to the frame along the hinge of the stem. First, we estimated the axis of the stem by solving a least-squares problem based on a calibration movement where the subject rotated the handlebars through the full range of motion when the bike was in a static position [21]. This method relies on the assumption that the change in orientation between the steer and the frame mainly occurs around one axis (i.e., the axis of the stem) during the calibration movement. Second, we used euler angles to compute the steering angle as the orientation of the stem with respect to the handlebars along the axis of the stem for all recorded frames.

The orientation of the sensors and the relative angles between sensors were analyzed along the vertical (gravity) axis by converting the rotation matrices to euler angles. The range of motion, defined as the maximal angle between sensors, was computed for the rotation of the pelvis with respect to the frame, the torso with respect to the frame and the torso with respect to the pelvis (Figure 1). The start and end of the rotation over the right and left shoulder was identified manually based on the angular velocity of the torso sensor along the vertical axis (Figure 1C). Task duration was calculated as the time between these manually determined start and end points.

### 2.4 Excluded data

To explore if young and older subjects used different kinematic strategies during this task we evaluated the range of motion between the IMU sensors on frame, pelvis and torso for the subjects that performed the trial successfully. The data of subjects that lost balance (i.e. foot on the ground, cycling outside the lane) or made mistakes in identifying the cones was excluded from the kinematic analysis. We additionally had to exclude data for several young and older subjects due to a faulty wireless connection between the IMU sensors and the base station. There was mainly a data loss for the sensor on the pelvis (17 young subjects, 9 older subjects) and to a lesser extent for sensors on frame, torso and handlebars (4 young and 4 older subjects). We therefore have a different number of participants depending if the outcome measure included the pelvis sensor or not (See Figure 2B and 3). Note that the errors in the task-performance were reported for all subjects (Figure 2A).

**Figure 2:**
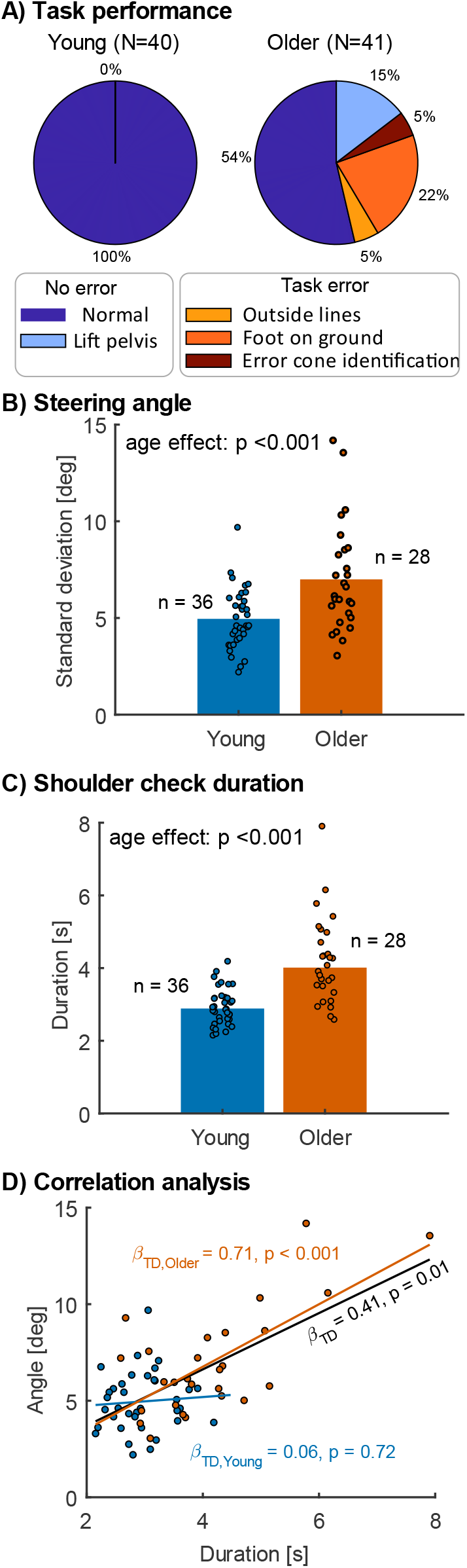
Task performance. A. 100% and 69% of the trials performed respectively by the young and older adults were successful (no error). B. The standard deviation in steering angle was lower in the young compared to the older subjects. C. The time needed to perform the shoulder check task was shorter in the young compared to the older subjects. D. The time needed to perform the shoulder check task was positively correlated with the standard deviation in steering angle across all subjects. When analyzed young and older adults separately, the association was not significant in the younger group, whereas it was strongly positive in the older group. TD, task duration.

### 2.5 Outcomes and statistics

The assumption of a normal distribution was tested using a Shapiro-wilk test for the data of the young and older subjects. Depending if the assumption of normal distribution was violated, we used a two-tailed Mann-Whitney U test or an independent t-test (ttest2, Matlab 2019) with an alpha of 0.05 to compare the relative range of motion of the sensors on the frame, pelvis and torso, the standard deviation in steering angle and the task duration between the young and older subjects. Additionally, we used robust linear regression (robustfit in MATLAB) to examine the relationship between task duration and the standard deviation in steering angle, while controlling for the effect of age group, following the approach of Vandevoorde and Orban de Xivry [22]. We also performed the robust linear regression for each age group separately to obtain group-specific standardized *β*-coefficients (*β*_TD_).

## 3. Results

Older adults had more difficulties than young adults in identifying the color of a cone behind them by looking over their shoulder. Even when older adults were successful in identifying the color of the cone, they more often made other task-related errors and used a different kinematic strategy than young adults.

### 3.1 Increased number of task-related errors in older adults

The shoulder check task was successful in 100% and 69% of the trials performed respectively by the young and older adults (Figure 2). For the older adults, placing one or both feet on the ground was observed as the most common error during the shoulder check task (22%), followed by cycling outside the lines (5%), and errors in identifying the color of the cone (5%). In addition, 15% of the healthy older adults lifted their pelvis from the saddle during the shoulder check task while all younger subjects remained seated.

Even when older adults performed the shoulder check successfully, they required more steering corrections than younger adults to remain stable and to continue cycling straight ahead while performing a shoulder check. We found that the standard deviation in steering angle in successful trials was larger in young compared to older subjects (Figure 2B, t(62) = -3.73, p < 0.001, Cohen’s d = 0.95). Furthermore, older adults took longer to perform the shoulder check task compared to younger participants (Figure 2C, t(62) = -5.75, p < 0.001, Cohen’s d = -1.46). This longer task duration, meaning longer time performing two competing motor tasks (rotate trunk and head to perceive environment vs. keep balance and direction), was positively associated with greater steering angle variability (Figure 2D, *β*_TD_ = 0.41, p = 0.01, r = 0.64). Furthermore, there was a significant interaction between task duration and age group *β*_I_ = 0.33, p = 0.03), indicating that the strength of this association differed between age groups. When analyzed separately, the association was not significant in the younger group (*β*_TD,Young_ = 0.06, p = 0.72), whereas it was strongly positive in the older group (*β*_TD,Old_ = 0.71, p < 0.001).

### 3.2 Increased torso rotation in the older adults

Older adults rotated significantly further with the torso with respect to the frame during the task than younger adults (Figure 3A, t(62) = -6.28, p < 0.001, Cohen’s d = -1.59). The increased rotation of the torso with respect to the frame was caused by an increased rotation of the pelvis with respect to the frame (Figure 3B, t(44) = -6.75, p < 0.001, Cohen’s d = -2.04) as we did not find any evidence for an age-related increase in the rotation of the torso with respect to the pelvis (Figure 3C, t(44) = 1.50, p = 0.14, Cohen’s d = -0.45). As a result, older subjects required less rotation in the neck and/or smaller eye-head rotations than young adults to identify the color of the cone (Figure 3D).

**Figure 3:**
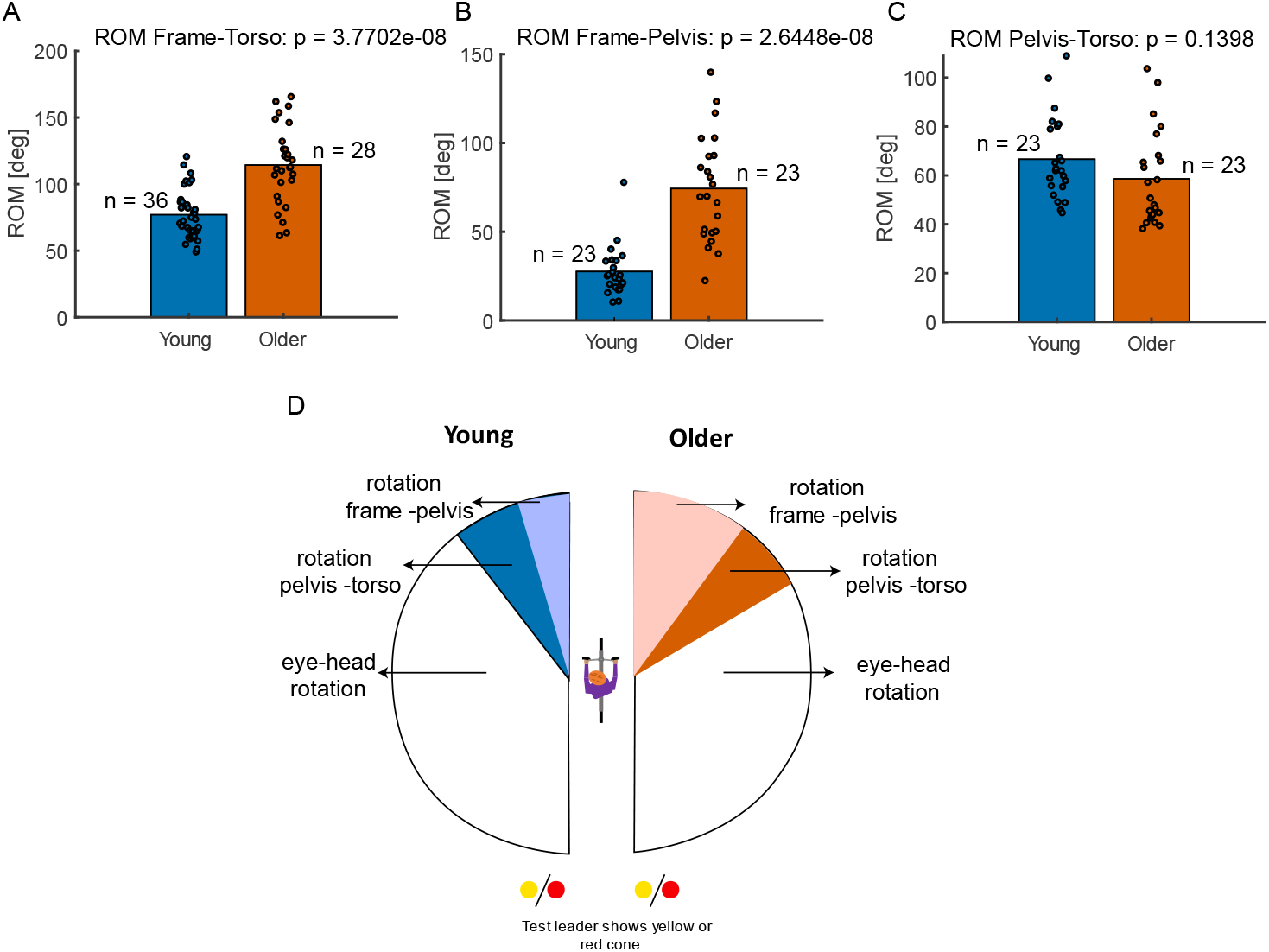
Results kinematic analysis. A. Older subjects rotated more with the torso with respect to the frame than young adults during the shoulder check. B. Older adults rotated more with the pelvis with respect to the frame than young adults during the shoulder check. C. The range of motion of the trunk with respect to the pelvis was not different in young and older adults. D. The average differences in kinematics between young and older adults is illustrated in pane D. Differences in torso-frame rotations imply that young adults relied more on rotation in the neck and/or eye-head rotation to identify the color of the cones than older subjects. Note that we excluded data of several young on older subjects due to (1) an error in the task or (2) a faulty wireless connection between the IMU sensors on the handlebars, frame, pelvis and torso during the task (see details in method section).

## 4. Discussion

Older cyclists performed a shoulder check in a worse manner than young cyclists. We asked participants to identify the color of objects behind them while controlling cycling direction. Even though our group of older cyclists was by no means representative for the oldest cyclists on the road as they were aged up to 65 years only, one third of them failed the task by either making cone-identification errors, interrupting cycling to prevent loss of balance, or cycling outside the lane while all young adults performed the task successfully (Fig. 2A). We also found that even when older subjects could perform the task successfully, they had more difficulties in performing the concurrent tasks of object identification and controlling steering direction, as the increase in variation in steering angle in the older subjects shows that they had to make more corrections to control the cycling direction or to control balance. These results are consistent with the study of Kovácsová et al. [20]. Unlike that study, our study is the first to investigate differences in kinematic strategies between younger and older participants when performing the shoulder check.

A first explanation for the increase in steering corrections is the different kinematic strategy used by the older subjects. Older subjects rotated more with the torso with respect to the frame (Fig. 3), possibly to compensate for a decreased axial rotation range of motion in the neck and in eye-head rotation [23]. This is in agreement with research in car drivers where a reduced range of motion in the neck, and functional neck range of motion during the task, in older adults was associated with an increase in target detection errors during a shoulder check [24]. Yet, in contrast to our results in biking, older participants exhibited less trunk rotation to check the blind spot than younger participants when driving a car [23], which directly impacted their ability to detect a target. Reduced neck range of motion has also been identified as a risk factor for older car drivers [25] but this can be improved by training neck, shoulder and trunk flexibility [26,27].

Alternatively, given the mechanical coupling between the trunk and the arms, the increased trunk rotation in the older cyclists might cause undesired arm motion, which results in undesired steering. Additional steering corrections are most likely needed to compensate for the co-rotation of the trunk and arms to control balance and cycling direction. Note that the neck and head-eye rotations strategy used by the young subjects is mechanically not directly linked to arm motion and therefore undesired steering angles. Hence, the difference in kinematic strategies (increase in torso rotation) during the shoulder check task might explain the increase in steering angle variance, and potentially also the increase in balance loss during the task in the older subjects. Several studies found that inter-joint coordination was not impaired in older people, suggesting that older people can compensate for the passive dynamics due to their own movements as well as younger people [28].

A third potential explanation for the increase in steering corrections is a reduced ability in older subjects to estimate the position of the arms in the absence of vision. When turning the head, the arm and the steer are located outside of the field of view. Given the age-related decrease in proprioceptive acuity [13,27–29, but see 30–32], older adults rely much more on vision than on proprioception to control movements such as bimanual upper limb movements [34], reaching or grasping [35,36] or walking [37]. The preponderance of visual inputs to maintain a stable steer control would explain why older adults exhibit more steer variability when the gaze is away. The plausibility of this alternative is reinforced by the fact that, during car driving, a multi-tasking condition that took the gaze away from the road led the older adults to exhibit deviations from their trajectories similar to those observed in the current study [38].

Finally, the increase in steering variability and decrease in task performance could also stem from the decrease in dual-tasking abilities with aging [17,39,40]. The shoulder check task requires participants to simultaneously bike straight and look behind. Our results showed that older adults performed this task more slowly than younger participants, likely due not only to slower motor planning but also to diminished dual-tasking capacity [40]. Paradoxically, taking more time on the task prolongs the period during which attention and gaze are diverted from the direction of travel, potentially compromising balance control. Indeed, our findings indicate that longer task duration is correlated with greater steering angle variability over all participants. Similar findings were seen during car driving, dual-tasking or multitasking made older participants drive in a slower and more variable way than their younger counterparts [38,41]. Dual tasking during driving causes an increase reaction time to brake, sloppy steer control and worse distance estimation, especially in older adults [38,42,43].

Our results raise awareness for balance threats imposed by shoulder checks in older cyclists, which can lead to deviations from the heading direction. These deviations increase the risk of encountering uneven terrain (e.g., curbs, soft borders) or colliding with surrounding objects, potentially resulting in accidents. To address this issue, a combination of infrastructure improvements and individual-level interventions should be considered. From an infrastructure perspective, policy makers could mitigate this risk by implementing wider cycling lanes, allowing cyclists to deviate from the direction of travel without endangering the cyclist. Additionally, prioritizing road designs that minimize the need for shoulder checks can reduce exposure to these balance threats (e.g. prioritize perpendicular crossings instead of merging lanes). On an individual level, cyclists may benefit from using assistive technologies like rear-view mirrors or radar systems. However, current devices often suffer from limited fields of view. Our results underscores the need for innovative, user-friendly solutions that improve rear visibility. Furthermore, targeted training programs to improve shoulder check technique, along with exercises to increase neck and trunk flexibility, may help older cyclists maintain better control and stability during these maneuvers. Flexibility training [44] or yoga [45] has been shown to improve spinal and shoulder mobility and could support shoulder check performance. However, there are currently no evidence-based programs specifically designed to improve shoulder checks. Further research is needed to identify effective training methods for older adults.

In conclusion, we demonstrated that the ability to perform a shoulder check task already drastically decreased in adults aged 60 to 65 as compared to young adults. As shoulder checks are a common task while cycling, e.g. to safely interact with other road users, the reduced ability to perform a shoulder check task while maintaining heading direction might contribute to the age-related increase in single-sided accidents [10]. While we need to further understand the causes, a combination of infrastructure improvements and individual-level interventions could counteract this age-related single-sided accidents.

## Conflict of interest

All authors hereby declare that there are no conflicts of interest.

## Competing interests

The author(s) declare no competing interests.

## Funding

Flemish agency for scientific research (FWO-Vlaanderen): FWO-12ZP120N

## Appendix 1 details on conventional bike and e-bike and sensor placement

The kinematics of the cyclist and bike was analyzed using inertial measurement units (IMU) mounted on the handlebars, frame, pelvis and torso (Figure appendix 1). Cycling data of all subjects was collected while riding the same conventional and electrical bicycles, provided by the KU Leuven. Both bicycles were similar models with step-through frames. Only a small difference in frame height was present, with the frame of the electrical bicycle being slightly higher. The weight of the electrical bicycle was 27.7 kg, which was 7.9 kg heavier than the conventional bicycle. The electrical bicycle had a front wheel hub motor, and the battery was located above the rear wheel. Power support could be set to five levels. In the present study the third level was chosen, which can be described as normal ‘support’. Each bicycle had six gears, which were set on the fourth level. Participants were asked to not change these settings during the whole experiment. Participants could change the saddle height to a comfortable height. Wearing a bicycle helmet was mandatory.

**Figure appendix 1:**
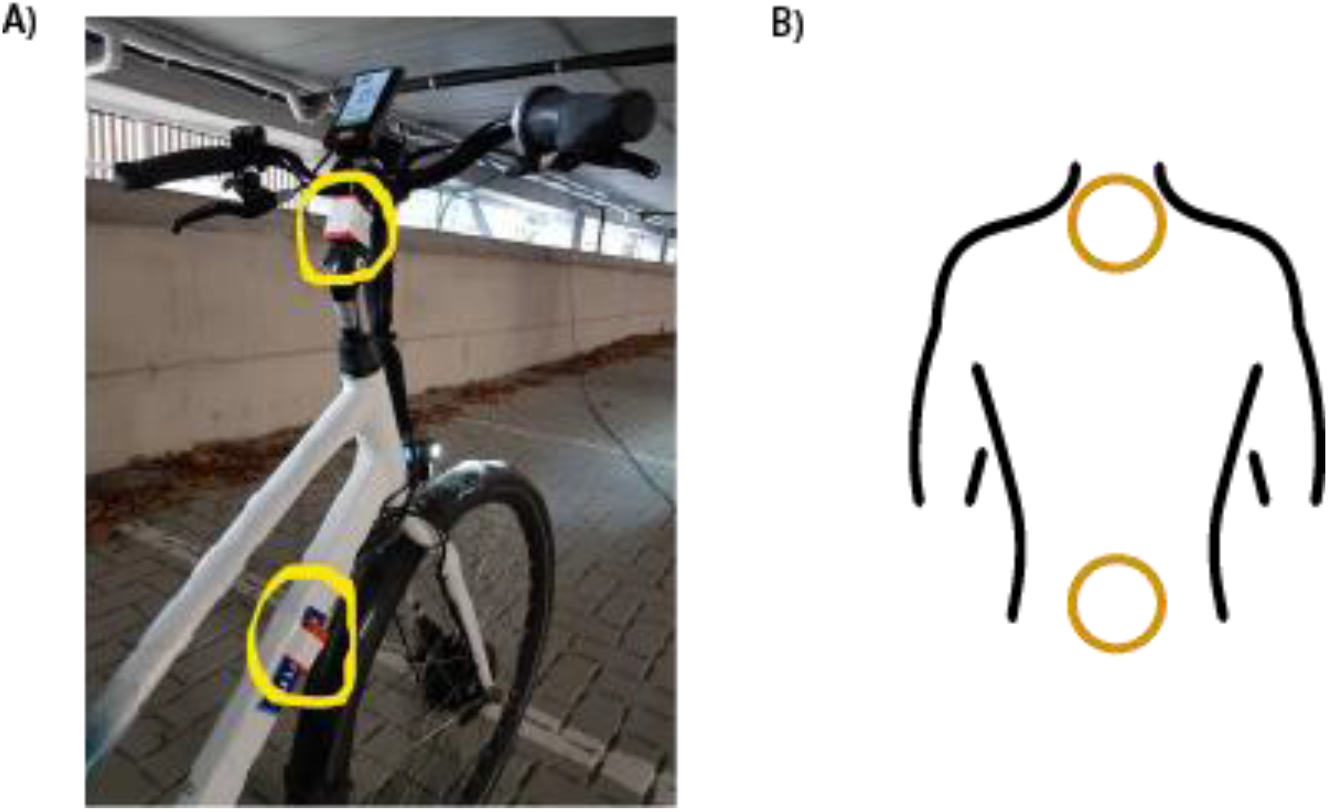
IMU placement on the frame and handlebars of the bike (A) and on the pelvis and torso of the subject (B)

## Appendix 2 Kinematic analysis for e-bike and conventional bike

We observed the difference between the young and older subjects in steering angle, ROM of the sensor, and task duration was similar on the conventional bike and e-bike during cycling in self-selected and slow speed conditions (Figure appendix 2).

**Figure appendix 2:**
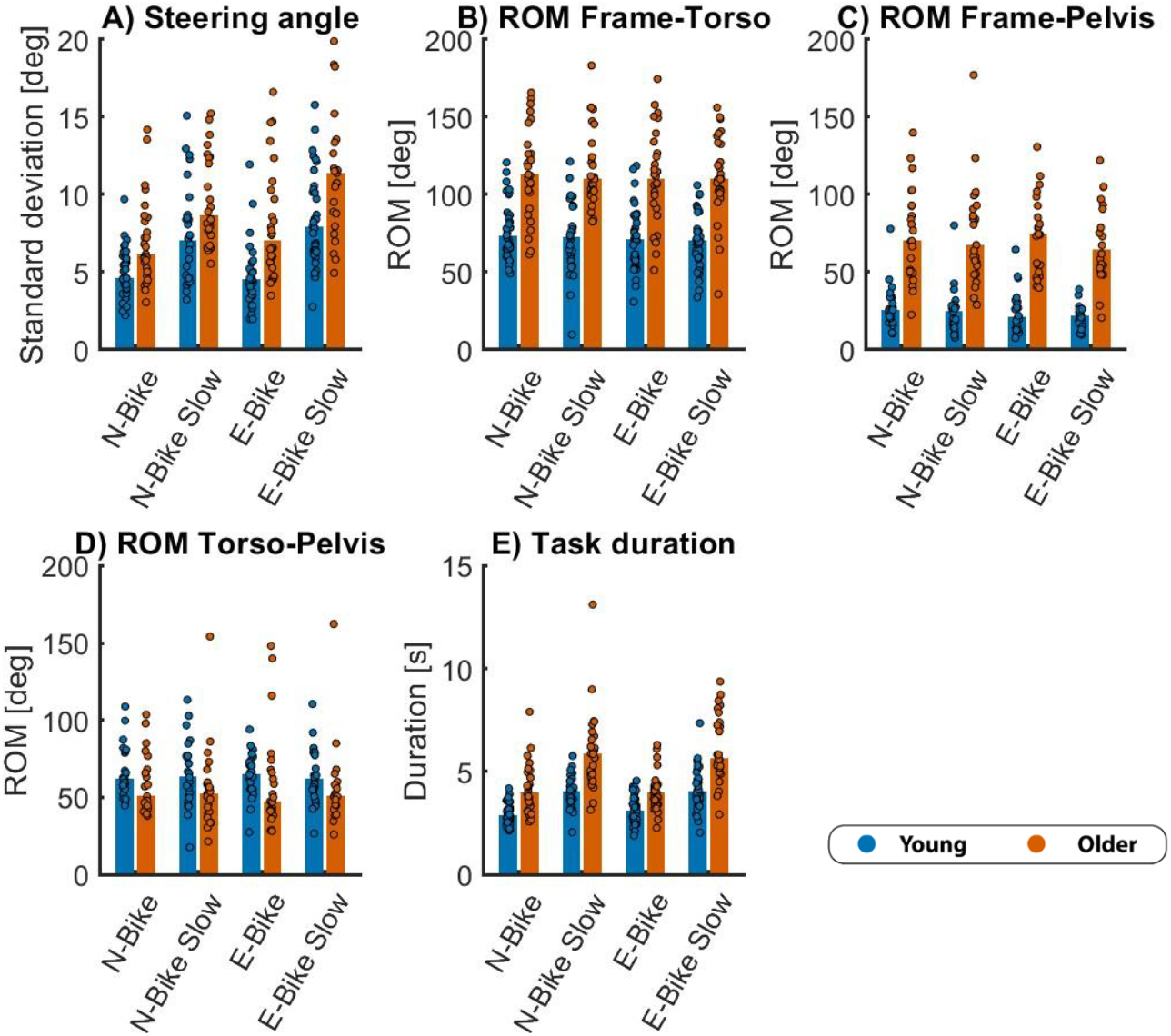
Standard deviation in steering angle (A) and range of motion between frame and torso (B), frame and pelvis (C), torso and pelvis (D), and task duration (E) during the shoulder check task when cycling with a conventional bike (N-Bike) at preferred or slow (N-Bike Slow) speed or with an e-bike (E-Bike) at preferred or slow speed (E-Bike Slow). We observed that the effect of age on steering angle, ROM, and task duration was similar for the conventional bike and e-bike at preferred and slower speed.

## Notes

### Competing Interest Statement

The authors have declared no competing interest.

### Summary of Updates

We have updated the manuscript with an additional analyses (Fig.2C and D) and with a more complete discussion about the solution for this age-related problem.

